# “Development and Implementation of Novel Chatbot-based Genomic Research Consent”

**DOI:** 10.1101/2023.01.23.525221

**Authors:** Erica D. Smith, Sarah K. Savage, E. Hallie Andrew, Gloria Mas Martin, Amanda H. Kahn-Kirby, Jonathan LoTempio, Emmanuèle Délot, Andrea J. Cohen, Georgia Pitsava, Seth Berger, Vincent A. Fusaro, Eric Vilain

## Abstract

**Objective:** To conduct a retrospective analysis comparing traditional human-based consenting to an automated chat-based consenting process.

**Materials and Methods:** We developed a new chat-based consent using our IRB-approved consent forms. We leveraged a previously developed platform (Gia®, or “Genetic Information Assistant”) to deliver the chat content to candidate participants. The content included information about the study, educational information, and a quiz to assess understanding. We analyzed 144 families referred to our study during a 6-month time period. A total of 37 families completed consent using the traditional process, while 35 families completed consent using Gia.

**Results:** Engagement rates were similar between both consenting methods. The median length of the consent conversation was shorter for Gia users compared to traditional (44 vs. 76 minutes). Additionally, the total time from referral to consent completion was faster with Gia (5 vs. 16 days). Within Gia, understanding was assessed with a 10-question quiz that most participants (96%) passed. Feedback about the chat consent indicated that 86% of participants had a positive experience.

**Discussion:** Using Gia resulted in time savings for both the participant and study staff. The chatbot enables studies to reach more potential candidates. We identified five key features related to human-centered design for developing a consent chat.

**Conclusion:** This analysis suggests that it is feasible to use an automated chatbot to scale obtaining informed consent for a genomics research study. We further identify a number of advantages when using a chatbot.

## BACKGROUND AND SIGNIFICANCE

Informed consent for most research studies is a complex manual process designed to meet the requirements of human subjects research regulations by providing potential enrollees with appropriate education and decisional autonomy. Historically, accomplishing this human-centered approach requires a substantial time commitment from study team members. Consent conversations are challenging due to coordination and scheduling logistics and the large amount of highly complex information that must be communicated to participants without overwhelming them.[1–3] Additionally, despite efforts to optimize the traditional consent process, reports have revealed poor understanding by some participants in genomic research studies, including 40% of participants not realizing that they are even enrolled in a study.[4]

Traditionally, informed consent is obtained via direct interaction (i.e., in-person, phone, or video call) between a member of the study staff (e.g., geneticist, genetic counselor, or research coordinator) and the individual considering enrollment or their legally authorized representative or guardian. This requires that the individual has geographic proximity or technological access to the study staff, all parties have the available time and resources to meet, and the study team reviews consent materials fully with potential subjects to ensure their comprehension of all concepts and implications. Consent conversations for either clinical or research genetic testing generally address results of uncertain significance, secondary genetic findings (both actionable and non-actionable), incidental findings (including the potential identification of unexpected familial relationships), risks relating to sharing genomic information, and potential sensitivities and fears around privacy and medical research.[5,6] Furthermore, some communities may face significant hurdles to research participation due to limited access to basic medical and research resources, as well as mistrust of research due to historical mistreatment.[7] Socio-cultural and language barriers may also hinder recruitment and obtaining informed consent.[8] Obtaining uniform informed consent is particularly important for equitable access to research opportunities and is particularly challenging in studies that require large-scale enrollment to achieve appropriate statistical power, even more so if children or vulnerable groups are involved.

Chatbots are computer programs designed to simulate human conversation. In healthcare, chatbots are used to screen for risk of hereditary breast and ovarian cancer,[9] colorectal cancer,[10,11] for vaccine management,[12] and even for mental health therapy,[13,14] among other health and disease contexts. They accomplish a variety of tasks, including collecting medical information, providing genetic information, returning test results, obtaining limited consent, and facilitating testing referrals to other family members in clinical settings.[15–17] Our study team worked with the Genetic Information Assistant (Gia®) chat software to develop a chat enabling a streamlined, auditable consent process for the Pediatric Mendelian Genomic Research Consortium (PMGRC), part of the national-scale Genomics Research to Elucidate the Genetics of Rare Diseases (GREGoR) consortium funded by NHGRI.[17] Though chatbots have been implemented successfully in a variety of clinical settings, to our knowledge this is the first analysis of using chatbot technology to facilitate complex informed consent for enrollment in a genomics research study.

## OBJECTIVE

We conducted a retrospective analysis of the implementation of a chat-based consent process, comparing the efficacy of this approach to traditional consent methods. Using principles of human-centered design, we identified five key factors relevant to developing a chatbot for informed consent. First, **understand the user’s context:** this includes understanding the individual’s motivations for participating in the research study and possible fears or concerns about research participation. The chat should adapt based on the user’s level of familiarity with the study and background information. Second, **make the consent process clear and easy to understand:** the chat should be designed to clearly explain the nature of the study, the risks and benefits of participating, and the user’s rights as a participant. The chatbot should also make it easy for the user to ask questions and get clarification on any confusing points.[18] Third, **respect the user’s agency:** the chat should fully describe the relevant considerations and allow flexibility around how and when individuals want to engage with the consent conversation. Fourth, **make the interaction as efficient as possible for the user:** the chatbot should enable users to complete the consent quickly and with minimal effort, saving time and providing a better experience. Fifth, **satisfy all regulatory and security requirements:** it is critical that the chat service meets security and regulatory requirements to protect the safety and privacy of users, and confidentiality of data. The chatbot should be Health Insurance Portability and Accountability Act (HIPAA) compliant and follow security best practices such as data encryption, user authentication, and limited access controls.

## MATERIALS AND METHODS

### Cohort design

We conducted a retrospective analysis of our Gia consent process compared to traditional consent methods (Figure 1). We analyzed the consent interactions and outcomes for all families who expressed interest in consenting to the PMGRC study over a period of six months, starting from the launch of the Gia option in June 2022. At the time of referral, each family could decide if they would prefer traditional consent with a study team member (in-person or via video call) or to use the chat-based consent process. Families were considered eligible for the chat-based consent if they were eligible for the PMGRC study and English-speaking. A total of 144 referrals for candidate families were included in this analysis, including 74 families who opted for traditional consent and 54 who elected to use the chat-based consent process. From this group of 144 referred families, 72 families comprising 109 adults had completed informed consent at the time of data collection. In this analysis, we focused on the adults who provided consent, not the 66 individuals for whom consent was obtained from their legal guardians. Demographic information for each participant was requested using a REDCap intake form after they consented to the study.

**Figure 1.**
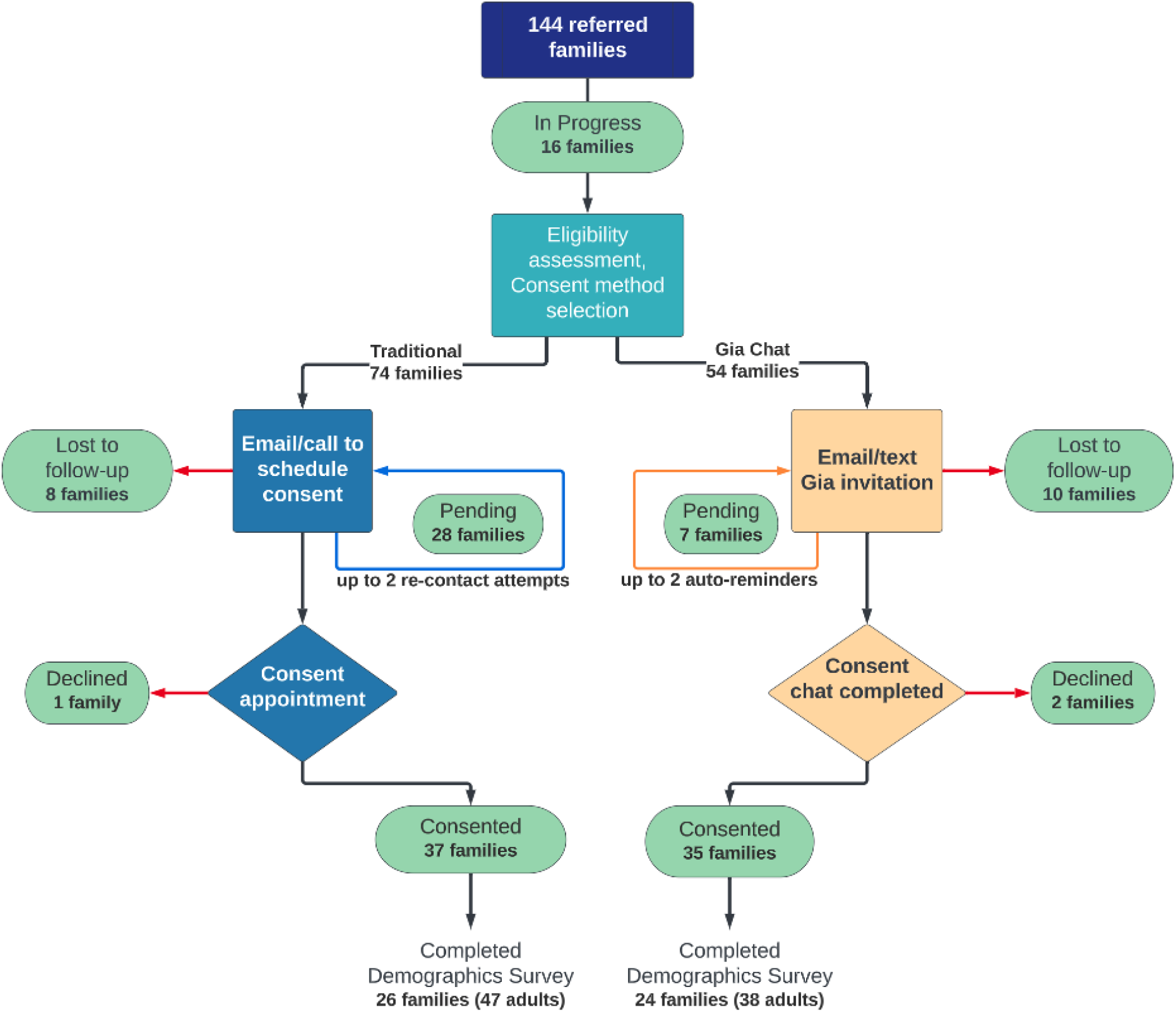
Consent status for referred families at the time of data collection. A flowchart showing the progression from referral to completion of consent and the demographics survey. Green ovals indicate the current status of referred families. This analysis includes the first 6 months during which Gia was offered for providing consent, June-December 2022.

### Gia chatbot platform

The chat-based consent experience was developed using Genetic Information Assistant (Gia ®), a HIPAA-compliant software platform developed by Invitae. Gia presents scripted content and allows users to choose among pre-populated responses at various intervals (Figure 2). The Gia technology supports a flexible user experience, with some mandatory content presented to all users and branching logic that presents optional content only when specific information is requested. Because the chats are web-based, private health data is not stored on the user’s device. Patients and families can securely interact with the chat on any web-connected device (e.g., smartphone, tablet, or computer) at their own convenience. Each user’s conversation progress is saved within their individual chat encounter, so individuals can start the chat and then come back later if necessary. The Gia interface simulates human conversation via text and is designed to be engaging, empathetic, and upbeat, qualities that are shown to increase user engagement and communication efficacy.[14]

**Figure 2.**
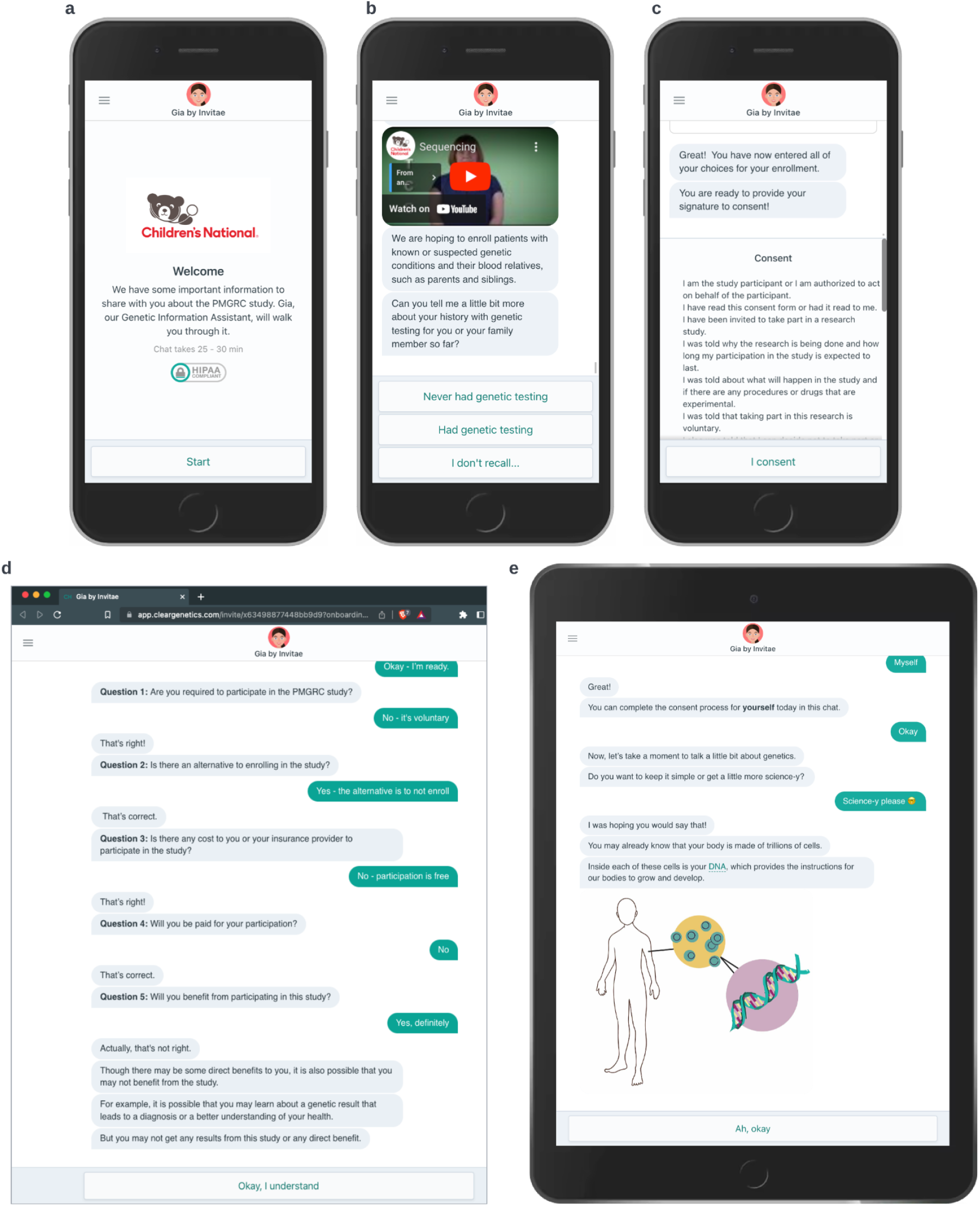
Examples of Gia chat interactions and accessibility. (A - C) Chat using a mobile device and highlights HIPAA compliance, educational video link outs, and the final consent. (D) Chat using a desktop web browser and the built-in teach back for incorrect quiz answers. (E) Chat using a handheld tablet showing the ability to select more in-depth information about genetics.

### Consent chat content

The Gia consent chat was developed by the study team based on the content of the existing IRB-approved research consent form used by study team members at Children’s National Hospital. The chat presents information at an eighth-grade reading level (Flesch-Kincaid Grade Level = 8, Flesch Reading Ease = 62.1). At the start of the conversation, Gia provides a brief orientation on how to use the interface and offers the alternative of consenting with a study team member. The initial sections cover general genetics information, benefits and risks of participation, privacy, and data sharing. Prospective participants are given the option to review either a basic or more detailed version of the genetics section.

Gia explains the types of research results that may be identified and enables users to select which types they would want to receive (primary results, actionable secondary results, and not actionable secondary results). It also explains that incidental results (results that are unexpected and unrelated to the reason for testing) will not be communicated to participants.

Prior to providing a consent signature, potential study participants or their legal guardian/representative must demonstrate an understanding of study participation by passing a knowledge assessment. The quiz has 10 questions based on an IRB-approved short form. All questions must be answered correctly before the individual can finalize their consent. Answering any question incorrectly results in a teach-back response and progression to the next question for the continuation of the quiz attempt. Users receive two attempts to pass the quiz. If they do not pass after the second attempt, they are directed to speak with a study team member.

The chat facilitates contact with study team members when additional support is needed, for example, when the individual has further questions or when pediatric assent is needed.

### Traditional consent

For candidates who opted to use traditional consent, the study team members made up to three attempts to schedule this conversation (typically via two emails and one phone call). Once the consent conversation was scheduled, the study team member sent an invitation to a personalized video call or scheduled an in-person visit. At the time of scheduled consent, the study team member and candidate participant reviewed the IRB-approved research consent form. For trios or larger families, the consent conversation could include multiple family members or separate conversations with each adult candidate participant.

### Gia consent

For candidates opting to use Gia, the study team entered relevant information into the Gia system, which then sent a unique web link to an individualized consent chat to the candidate participant or their parent/guardian(s). This link was delivered via email or SMS text, depending on their expressed preference. If the individual did not complete the chat, the chat system automatically provided up to two reminders to engage with the chat, at 2 weeks and at 4 weeks. The chat was designed to allow adult probands and family members (18 years or older) to provide informed consent. For families, each adult family member who is considering enrollment must complete their own unique encounter with the chat. For individuals not capable of providing their own consent based on age or developmental status, guardians can complete consent conversations on their behalf. Those who proceed to the consent segment may provide consent for themselves and/or for up to three children in a single chat encounter.

### Assent for both traditional and Gia consent

For children who are of age to assent (7 - 17 years of age), parents may provide their consent, and then assent is completed with a study team member.

## RESULTS

### Description of the cohort

We compared the enrollment outcomes of 144 families referred over a six-month period between traditional and Gia consent (Figure 1). Our study is specifically attempting to recruit families in trios (i.e., mother, father, and proband). Using traditional consent, 8 families were lost to follow-up and 1 family declined, compared to Gia, where 10 families were lost to follow-up and 2 families declined. Ultimately, 37 families completed the consent using the traditional process, compared to 35 families using Gia, a result that was not significantly different *x*^2^(1, n=128) = 2.78, p=0.095. By contrast, we observed a significant difference in the number of families classified as “pending”: 28 families in the traditional process, compared to 7 families using Gia *x*^2^(1, n=128) = 9.7, p=0.0018 (Figure 3).

**Figure 3.**
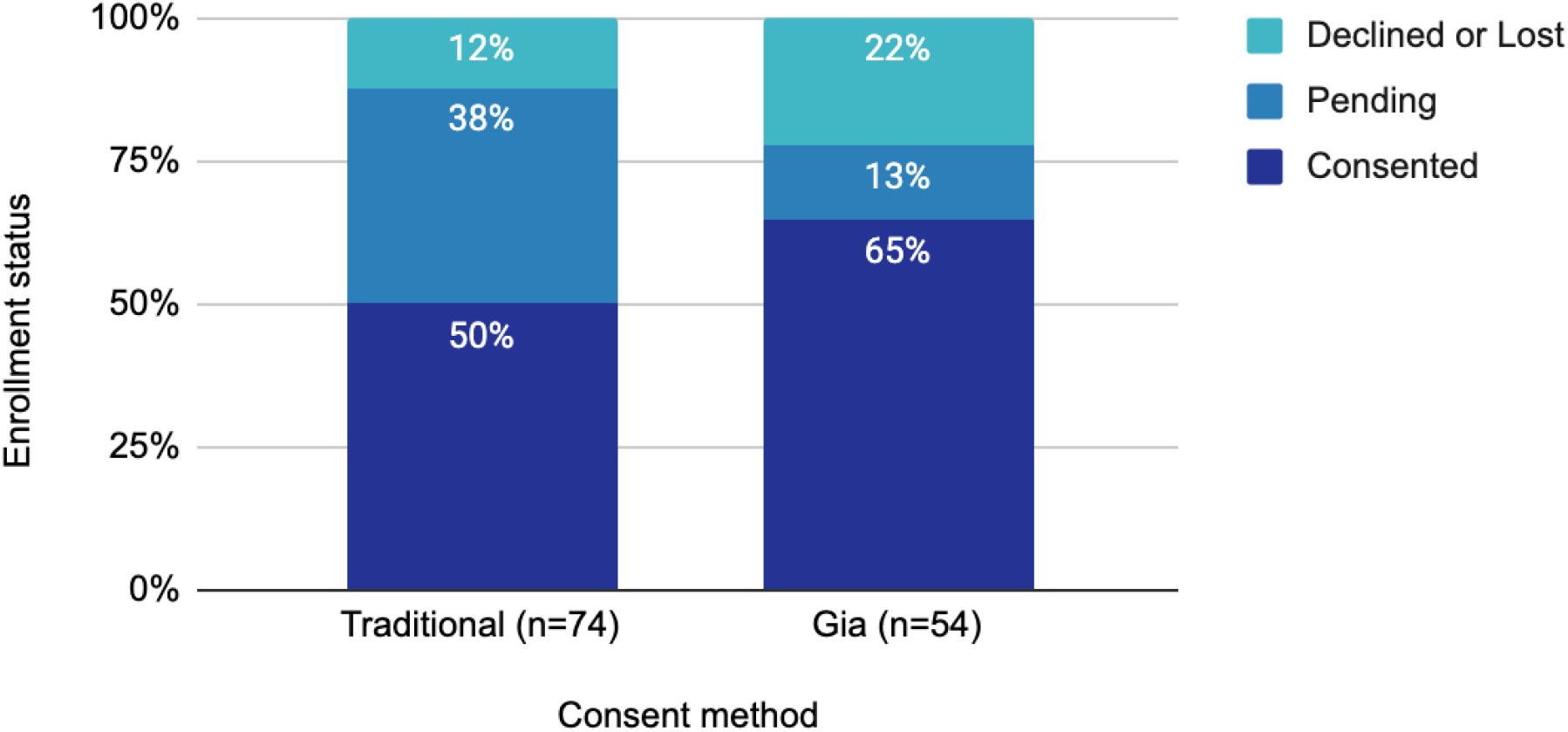
Consent outcomes for referred families. A representation of the proportion of referred families in each consent group who: 1) completed the consent and declined or are considered lost to follow-up (Gia - 12 families, Traditional - 9 families); 2) are pending and have not yet completed the consent process (Gia - 7 families, Traditional - 28 families); or 3) completed the consent to enroll (Gia - 35 families, Traditional - 37 families).

A total of 109 individuals consented to the study (59 traditional, 50 Gia). A survey was subsequently sent to all consented individuals, resulting in demographic information for 47 traditionally consented individuals and 38 who consented via Gia. Analysis of this information shows no significant differences between participants who chose each method. The median age of respondents is similar between Gia and traditional consent groups (p=0.84 by 2-way t-test). There was no significant effect of self-reported sex on consent method choice *x*^2^(1, n=83) = 0.52, p=0.47. Ethnic composition of the two groups was not different either, with the majority of participants in both groups self-identified as “White” (Table 1).

**Table 1:** Self-reported demographics of consenting participants.

| <u>Participant characteristics</u> |  | <u>Traditional</u> | <u>Gia</u> |
| --- | --- | --- | --- |
| Age, median years |  | 45 | 42.5 |
| Age, range in years |  | 18-69 | 27-72 |
| Self-reported sex, n (% of method) |  |  |  |
|  | Female | 25 (53) | 22 (61) |
|  | Male | 22 (47) | 14 (39) |
| Race/Ethnicity, n (% of method) |  |  |  |
|  | American Indian / Alaska Native | 2 (4.3) | 2 (5.3) |
|  | Asian | 0 | 2 (5.3) |
|  | Native Hawaiian or Other Pacific Islander | 0 | 0 |
|  | Black or African American | 2 (4.3) | 1 (2.6) |
|  | White | 42 (89.4) | 32 (84.2) |
|  | Middle Eastern or Northern African | 0 | 1 (2.6) |
|  | More than one race | 2 (4.3) | 3 (7.9) |
|  | Unknown | 0 | 0 |
|  | Hispanic | 5 (10.6) | 6 (15.8) |

### Analyzing consent and enrollment time

Participants who elected to use the traditional consent process met with study team members either in clinic or over video call. We estimated the duration of these consent conversations by reviewing video call records for 28 consented families (9 families consented in person). The traditional consent conversation took a median time of 76 minutes per encounter (range 43 - 134 minutes). It was most common for the study team member to meet with candidate participants individually, as in 64% of the video calls (n=18). However, in some cases, multiple family members engaged in the same consent video chat: 21% of calls had two candidate participants (n=6), 11% of the calls had three (n=3), and 4% had five (n=1).

Engagement patterns differed between participants using Gia compared to traditional consent. A total of 93 chat invitations were sent to individuals who requested to consent with Gia. The chat was started by 68 individuals and subsequently completed by 55 of them. The majority of chat users (62%, n=34) completed their consent conversation with Gia in less than one hour (Figure 4A). The median completion time was 44 minutes. This was significantly less than the median traditional consent time (p=0.0083, 2-tailed Mann-Whitney test). Many Gia users paused the conversation and returned to complete it later, and these breaks are included in the total completion time. A few users (7%, n=4) spread the consent conversation over multiple days.

**Figure 4:**
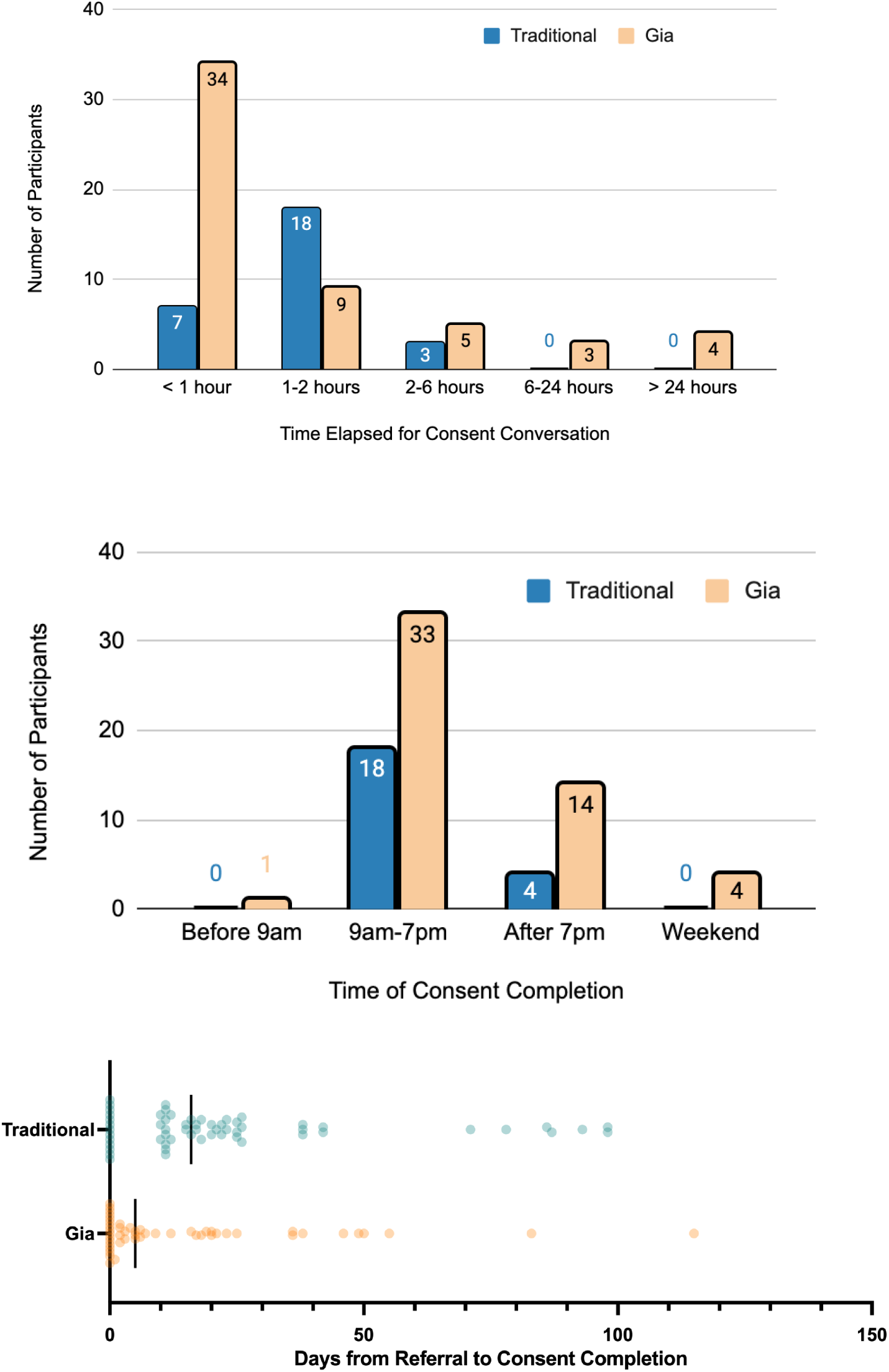
Gia engagement patterns. (A) Duration of consent conversation: Amount of time elapsed between opening Gia and completing the consent conversation. The time elapsed to completion includes breaks. For the Traditional method, time displayed is the duration of the uninterrupted video chat. (B) Proportion of participants who signed the consent form before 9am, between 9am - 7pm, after 7pm, and on a Saturday or a Sunday. (C) Distribution of the number of days elapsed between REDCap referral and consent completion for each consented adult. Using Gia allowed 16 subjects to enroll on the same day that they heard about the study. Traditional consent was used to enroll 13 subjects on the same day they learned of the study.

Additionally, participants frequently engaged in consent conversations outside of standard business hours. While 63% of Gia users signed the consent form between 9 am-7 pm on weekdays, the remainder signed the consent form during times when study team members are not accessible (Figure 4B). Notably, 27% of Gia users completed the conversation after 7 pm and 8% on weekends.

The total number of days needed to enroll subjects differed significantly between Gia and traditional methods. The Gia consent process was significantly associated with faster progression from referral to consent completion (Figure 4C). The median time from referral to consent was faster by 11 days using Gia compared to traditional consent (5 days for Gia vs. 16 days for traditional p=0.0222 by 2-tailed Mann Whitney test). Additionally, the option to offer consent via Gia enabled the study team to offer same-day consent and engage candidate participants who expressed interest during a family advocacy conference and during busy clinic days, when the study team would not have had the time to enroll these families using traditional methods alone.

### Assessing informed consent

Validation that participants understood key information is built into Gia (questions in Supplemental Information). Structured knowledge checks are not currently part of the standard of practice for traditional informed consent conversations, and therefore comparisons between the two consent mechanisms are not possible. However, the analysis of quiz performance allows for some unique insights regarding participants’ understanding of study participation. First, of the 59 individuals who took the quiz, understanding was generally high, with ~96% of users successfully passing. Most (76%) subjects passed the quiz on the first attempt, and 20% of patients passed on the second attempt. Second, the quiz accurately identified situations in which the user was not able to provide informed consent. Two users failed the quiz twice: one due to inebriation and a second user due to intellectual disability. In both cases, a team member followed up with the participant/legal guardian. All other Gia users completed their consent or decided to decline participation without any additional support or discussion with the study team.

Review of quiz responses shows possible gaps in understanding after chatting with Gia. The question most commonly answered incorrectly was “Will you benefit from participating in this study?”. The correct answer is “Maybe, but not necessarily,” but 3 individuals responded “No, definitely not,” while 4 responded “Yes”. Candidate participants had difficulty correctly answering the question “Are there any risks from participating?”, as 4 individuals responded “There are no possible risks”, while the correct answer is “Yes, there may be risks.”

### Evaluating user feedback

When given the option to choose the level of detail they wanted in the genetics education portion of the chat, 73% of Gia users chose the basic version, and 27% chose the more detailed version.

Finally, we gathered participant feedback about the chat consent experience. The final display of every completed consent conversation was the prompt “Rate your experience - We’d love to get feedback on Gia.” Responses were gathered using a Likert scale with three faces: happy, neutral, and sad. Out of 42 responses, 36 (86%) were positive and 6 (14%) were neutral. This suggests that most participants had a positive experience using the chat.

## DISCUSSION

In this study, we developed an efficient consent chat using a human-centered design approach while maintaining the necessary requirements for human subject research and informed consent. Our goal was to create a tool that performed as well as or better than the traditional human consent process to improve study scalability. Overall, we demonstrated that Gia performed similarly to traditional consent, which indicates that it is a viable alternative during the consenting process.

In terms of our approach, we addressed five aspects of human-centered design relevant to designing a consent chat. First, **understand the user’s context:** candidate participants were given the option to consent via traditional methods or using Gia to ensure that we could meet their accessibility needs. Gia operates in a regulated environment, meaning that some topics were required to be included in the chat, and any major changes needed to be approved by our IRB before release. Regardless of these constraints, the chat allows a flexible user experience. Some mandatory content is presented to all users and some optional content is only displayed to those users who request more detailed information, such as determining the level of detail of background genetic information, getting responses to common questions, or contacting a team member.

Second, **make the consent process clear and easy to understand.** We condensed a >7,000-word 13-page consent form into a script at an 8th grade reading level and presented it in chat form on a web-enabled device. We ensured Gia fully explained the study along with the risks and benefits of participating and the participant’s rights. This information was given in bite-size chunks representative of a chat conversation. Gia provides 3-4 sentences to the candidate participant and then pauses for user interaction responses such as “Okay” or “Good to know!”. The effect is to make the content more engaging and the pace more consistent with a human conversation, reducing user burnout or “click through.”

Using a chat-based consent also allowed us to confirm participant understanding through quizzes. This is an improvement over traditional consent methods, as passing the quiz confirms that consent is truly “informed.” This is particularly advantageous for genomics research consent, which requires participants to understand multiple complex concepts. Additionally, the ability to gather data about participant understanding also offers opportunities to further study consenting practices for research and process improvement. For instance, reviewing which questions are missed frequently will help the study team improve specific parts of the consent conversation.

Third, **respect the user’s agency.** The chat logic was written to allow users to determine how and when they wish to engage. The chat-based consent format allows flexible access, providing an asynchronous method to engage candidate participants at their convenience. This increases access to candidate participants who may not have time to consent during the workday. Engagement patterns showed that most chat users accessed and completed the informed consent conversation without study team member intervention. Many participants engaged with the chat on evenings and weekends or spread the chat over multiple sessions, patterns that are not available using traditional consent methods. This may be of particular value to candidate families who do not have the option to take time away from work during typical business hours, or who are not available simultaneously.

Fourth, **make the interaction as efficient as possible.** We observed significant time savings, both within the consent conversation as well as the time from referral to completed consent. This process improvement is reflected in the significantly lower rate of families still “Pending” in the Gia workflow, compared to the traditional consent. Regardless of the pathway of informed consent, a study team member was still essential to answer participant questions, review health records to determine eligibility, and obtain assent – all of which would have received less attention given the precedence that a traditional consent conversation takes in recruitment. Even though the entire consent process is not fully automated through use of Gia (e.g., assent requires speaking with a study member), implementing this method shifts the time demands of team members from routine conversations to focus on more complex tasks. Indeed our data show that shifting the burden from study staff to Gia for traditional consent conversations saved an hour or more per family. These results indicate that using a consent chat can save time for participants, allow study teams to dramatically scale study enrollment, and use team member time more efficiently.

Fifth, **satisfy all regulatory and security requirements.** It is paramount to human-centered design that the chat system we designed is compliant with regulatory requirements and security best practices to protect the safety, privacy of candidate participants and confidentiality of data. Gia is HIPAA-compliant, and access controls prevent unauthorized access to sensitive information.

The current study highlighted several challenges remaining in the consent workflows, even after implementing changes with the chat-based consent. For instance, in both consent methods, it is difficult to convey to parents that the study requires consent for each individual in the trio. Using Gia, multiple participants completed consent for their affected child but not for themselves, although they were happy to do so when prompted. Frequently, one parent remains pending and requires follow-up from the study team, regardless of whether they intend to use Gia or speak with a study team member in traditional consent.

A limitation of using this study to compare enrollment outcomes is that candidate participants were not randomly assigned to their consent methods. It is possible that people self-sorted by *a priori* intent; for example, individuals with more questions and reservations may have preferred to speak with a study team member, while individuals comfortable with technology might prefer a chatbot. Randomizing a larger group of prospective candidates to consent methods might be a better test of any possible effect of the consent method on enrollment outcome, albeit at the expense of individual agency in consent choice. Also, the chat is currently only offered in English, and thus non-English speaking participants were routed through the traditional consent process with a translator and not included in this study. While translation is a target for future development, it was beyond the scope of this initial implementation. Finally, there are remaining opportunities for development, as the assent process is not fully accommodated by the chat. Currently, the chat is designed to obtain consent from individuals age 18 or older, and individuals age 7-17 must speak with a study team member to provide assent. Incorporating a process to provide pediatric assent within Gia is an important area for future development.

## CONCLUSION

In the current study, we demonstrated that the consent completion rates are similar when using a chatbot compared to traditional consent. In addition, we identified five human-centered design features essential for a successful chat experience. We further identify a number of advantages when using the chatbot, such as a more flexible consenting experience for participants, faster time to consent, less time burden on study staff (resulting in the ability to recruit larger cohorts), and testing the understanding of participants.

While the analysis focused on the use of a chatbot in the setting of a genomic research study, it is likely that this consent approach could offer similar value in other research and clinical settings. Consent for clinical exome or genome analyses typically requires a substantial time commitment from members of the clinical team. Incorporating a chat-based clinical consent could expedite this process. Overall, this analysis demonstrates that the option to use a chat-based consent can streamline the consent process, allow more efficient use of study resources, and provide candidate participants with an effective and flexible consent experience.

## Acknowledgments

The authors would like to thank all families who participated in this research.

## Ethics, Funding, and data management

The Pediatric Mendelian Genomics Research Center (PGMRC) is a collaboration of Children’s National Research Institute, Invitae Corporation, and University of California Irvine, funded by the National Human Genome Research Institute (NHGRI) grant 1U01HG011745, as part of the GREGoR Consortium. This work was conducted under Children’s National IRB-approved protocol for the PMGRC study (Pro00015852). Data were collected and managed using REDCap electronic data capture tools hosted at Children’s National Hospital.[19,20] REDCap is a secure, web-based software platform designed to support data capture for research studies.

## Supplemental information

### Demonstration Gia link

https://app.cleargenetics.com/invite/x63498877448bb9d9?onboarding&demo&noresume

### Gia Quiz content

**Gia Quiz content:**
| Question | Correct Answer | Frequency of incorrect response (on first attempt) |
| --- | --- | --- |
| Question 1: Are you required to participate in the PMGRC study? | No - it's voluntary | 2/59 |
| Question 2: Is there an alternative to enrolling in the study? | Yes - the alternative is to not enroll | 2/59 |
| Question 3: Is there any cost to you or your insurance provider to participate in the study? | No - participation is free | 1/59 |
| Question 4: Will you be paid for your participation? | No | 0/59 |
| Question 5: Will you benefit from participating in this study? | Maybe, but not necessarily | 7/59 |
| Question 6: Are there any risks from participating? | Yes, there may be risks | 4/59 |
| Question 7: What is the purpose of the study? | To discover the causes of genetic health issues and better understand the role of genetic variants | 1/59 |
| Question 8: Will you need to provide samples and/or data? | Yes, the study team may collect some samples and/or data. | 0/59 |
| Question 9: Will your personal identifiable data be kept confidential? | Yes, the study team will work to keep my information confidential | 1/59 |
| Question 10: If you are injured as part of your study participation, what can you do? | I can contact the principal investigator | 1/59 |

